# Brain structure in pediatric Tourette syndrome

**DOI:** 10.1101/054437

**Authors:** Alton C. Williams, Deanna J. Greene, Jonathan M. Koller, Bradley L. Schlaggar, Kevin J. Black, The Tourette Association of America Neuroimaging Consortium

## Abstract

Previous studies of brain structure in Tourette syndrome (TS) have produced mixed results, and most had modest sample sizes. In the present multi-center study, we used structural MRI to compare 103 children and adolescents with TS to a well-matched group of 103 children without tics. We applied voxel-based morphometry methods to test gray matter (GM) and white matter (WM) volume differences between diagnostic groups, accounting for MRI scanner and sequence, age, sex and total GM + WM volume. The TS group demonstrated greater GM volume in posterior thalamus, hypothalamus and midbrain, and lower WM volume bilaterally in orbital and medial prefrontal cortex. These results demonstrate evidence for abnormal brain structure in children and youth with TS, consistent with and extending previous findings. As orbital cortex is reciprocally connected with hypothalamus, our results suggest that structural abnormalities in these regions may relate to abnormal behavioral inhibition and somatic monitoring in TS.

## Introduction

Tourette syndrome (TS) is a developmental disorder of the central nervous system defined by the chronic presence of primary motor and vocal tics (American Psychiatric Association, 2013). Tics are repeated, nonrhythmic, unwanted but usually suppressible movements or vocalizations (Black, 2010). TS usually involves one or more additional features, most often obsessions, compulsions, distractibility or impulsivity (Leckman et al., 2014). A clear neurobiological explanation for TS is not yet available, but research has provided many relevant clues (Mink, 2006; Martino and Leckman, 2013).

A number of studies have now examined the structure of the living brain in TS and have found significant changes in various brain regions compared to tic-free healthy control subjects (Greene et al., 2013; Church and Schlaggar, 2014). The largest studies were reported by Peterson and colleagues, with over 100 children and adults with TS and a similar number of control subjects. However, substantial questions remain about the structural anatomy of the brain in TS because methods and results have varied widely across studies, and because most studies were from small samples. A multi-center collaborative approach to brain imaging in TS might address these and other concerns.

Here we report the first analysis from such a collaboration, the Tourette Association of America Neuroimaging Consortium, applying structural MRI to large, well-matched groups of children and adolescents with and without TS.

## Results

### Subjects

Existing and newly acquired T1-weighted MP-RAGE images were collected from over 400 children age 7-17 years, including 230 with a chronic tic disorder (TS or Chronic Tic Disorder). After excluding scans with visible artifact, MPRAGE images were available from 109 TS subjects and 169 control subjects age 7-17 years. Of the 109 TS subjects, 103 could be matched one to one with a control subject for age (within 0.5 years), sex and handedness. Demographic and illness variables for these subjects are summarized in Table 1.

**Table 1.** Demographics and illness variables.

| **Table 1. Demographics and illness variables** |  |  |  |
| --- | --- | --- | --- |
|  | **TS group** | **Control group** | ***p*** |
| N, total → used | 230 → 103 | 216 → 103 | 1.00 |
| N by site, total → used |  |  | 0.000<sup>e</sup> |
| WUSTL | 141 → 78 | 156 → 70 |  |
| UCLA | 51 → 13 | 0 → 0 |  |
| NYU | 25 → 8 | 38 → 21 |  |
| KKI | 13 → 4 | 22 → 12 |  |
| Age (years, mean ± SD)<sup>a</sup> | 11.9 ± 2.1 | 11.9 ± 2.1 | 0.96 |
| Sex (M : F) | 81 : 22 | 81 : 22 | 1.00 |
| Handedness (# right-handed) | 103 | 103 | 1.00 |
| YGTSS Total Tic Score | 17.2 ± 8.6 (N = 64)<sup>b</sup> | n/a | — |
| ADHD clinical diagnosis | 55% (43 of 78)<sup>b</sup> | —<sup>d</sup> | — |
| CY-BOCS score (mean ± SD) | 5.3 ± 6.8 (N = 65)<sup>b</sup> | —<sup>d</sup> | — |
| OCD clinical diagnosis | 47% (15 of 32)<sup>b, c</sup> | —<sup>d</sup> | — |
| IQ (mean ± SD) | 108 ± 13 (N = 80)<sup>b</sup> | 119 ± 12 (N = 39)<sup>b</sup> | 0.000<sup>f</sup> |
| <sup>a</sup> Starting with this row, data describe only the final 206 subjects.<br><sup>b</sup> Not available for all subjects.<br><sup>c</sup> 45% (35 of 78) had CY-BOCS score > 0.<br><sup>d</sup> Not available for most control subjects.<br><sup>e</sup> $\chi^2 = 56.4$, 3 df.<br><sup>f</sup> $t = 4.53$, 81.9 df (two-sided *t* test, unequal variance, Welch df modification). | | | |

All images were T1-weighted 3D MPRAGE images, voxel size 0.9-1.8mm^3^, from different sites and different MR sequences. In all, 8 different scanner / sequence combinations were used to acquire the images (Supplemental Table 1).

**Supplemental Table 1.**
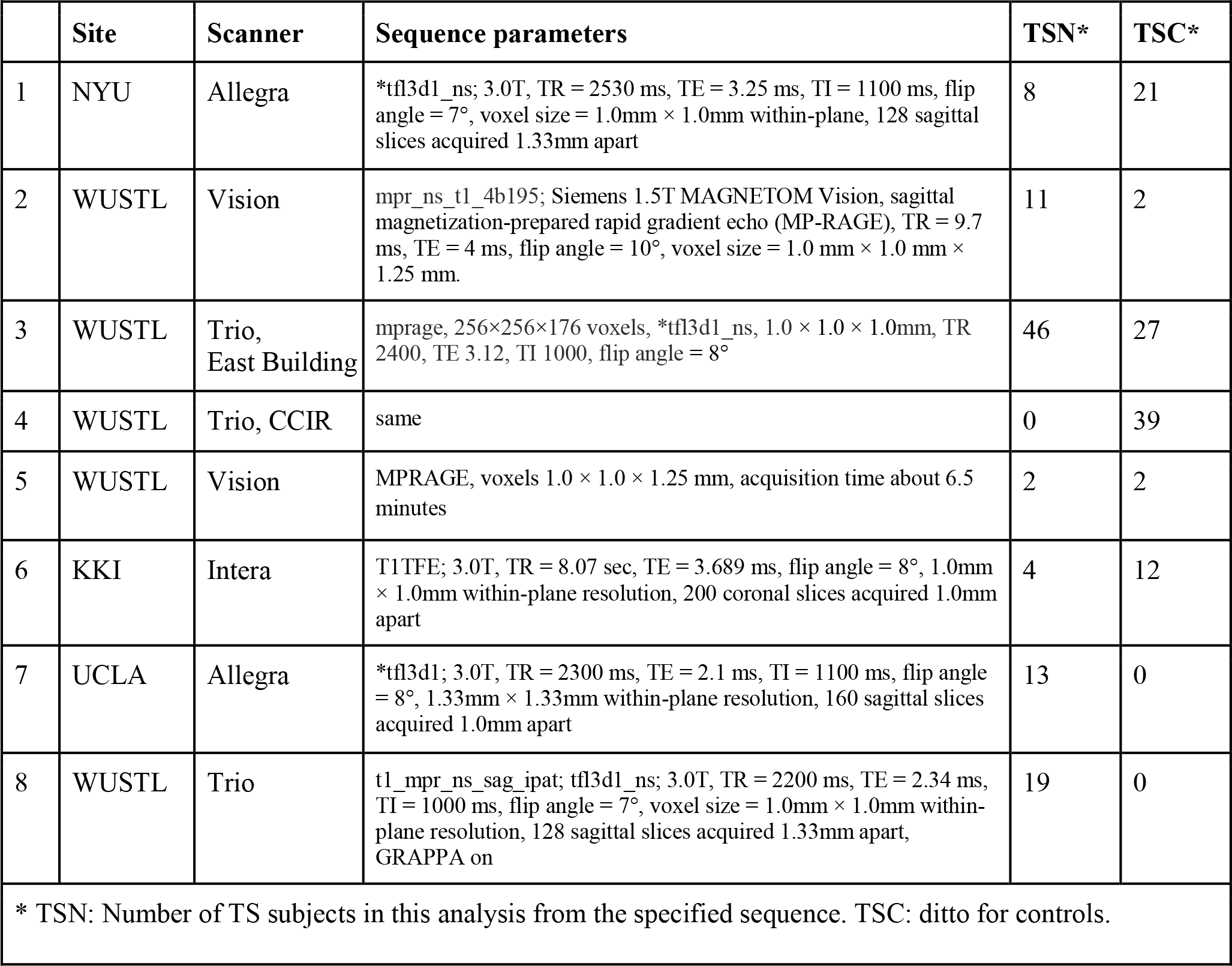
Image collection specifications

### Global volumes

Total GM volume was significantly correlated with age (*p* = 4.23 × 10^−7^), but no other factors, covariates or interactions were significant in the analysis of total GM or total WM (*p* ≥ 0.30).

### Regional differences in white matter volume in TS

Table 2 summarizes the VBM results for WM. Two fairly symmetric WM clusters were statistically significant after correction for multiple comparisons, both with lower WM volume in TS. The cluster volumes were 5.2 and 4.6ml, each corrected p = .001, and they were located in WM deep to orbital and medial prefrontal cortex (see Figure 1 and Supplemental Figure 1).

**Figure 1.**
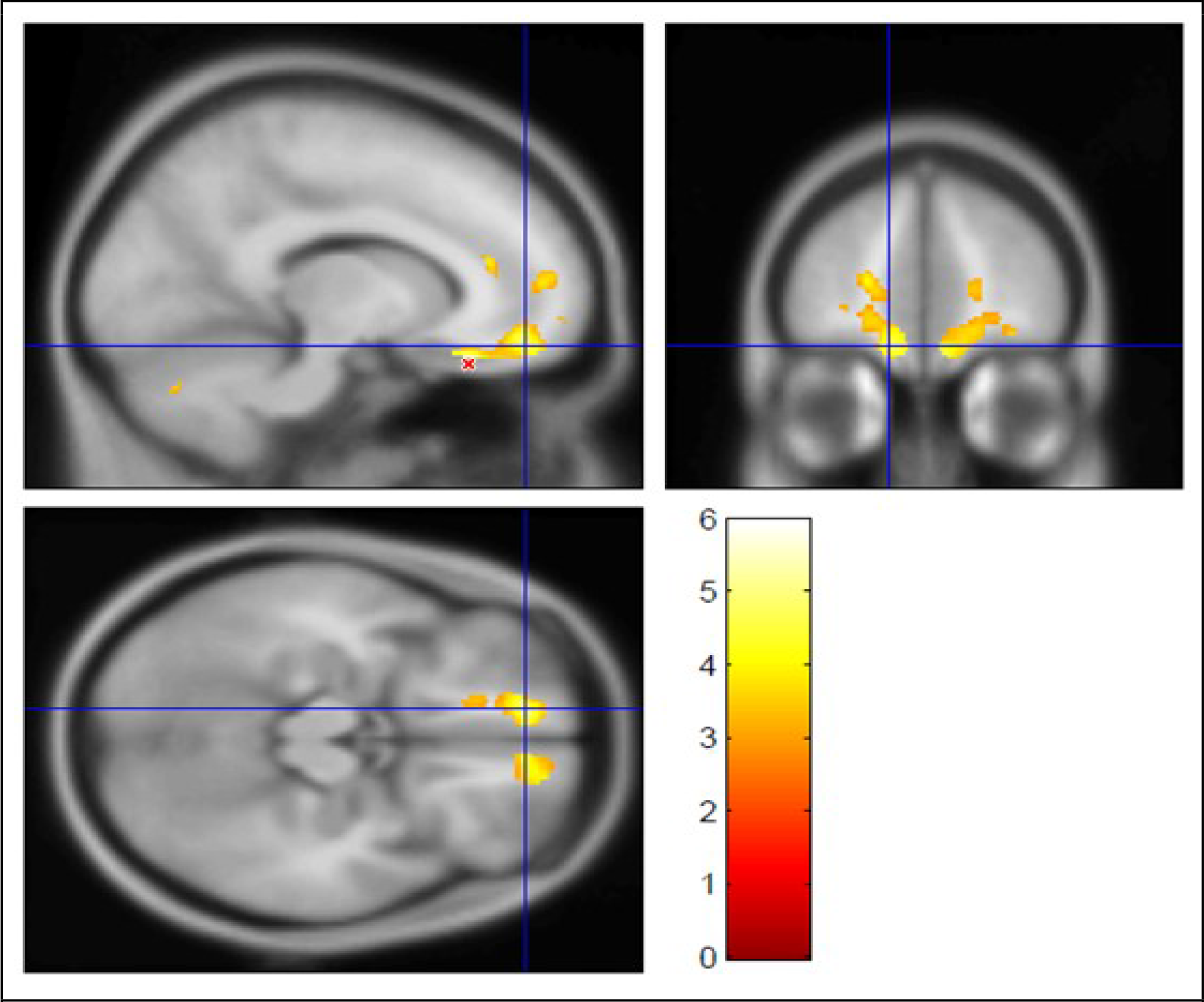
Largest cluster showing lower wm volume in TS compared to controls **Legend:** The largest cluster from the contrast showing where WM volume is lower in TS than in the control group (5.2 ml,*p*_FDR_ = .001; see Table 1). The *t* statistic is shown in color (thresholded at *t* ≥ 3.0), laid over the average MP-RAGE image from the entire sample (in grayscale). The crosshairs show (−12, 49.5, −16.5)_MNI_, left medial orbital gyrus, BA11. The peak *t* value from this contrast, t_193_ = 5.95, is at (−13.5,31.5, −22.5)_MNI_ in left medial orbital gyrus, BA13, near the red 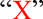 in the sagittal image. Supplemental Figure 1 shows the other significant cluster from this contrast, the homologous area on the right side of the brain.

**Table 2.**
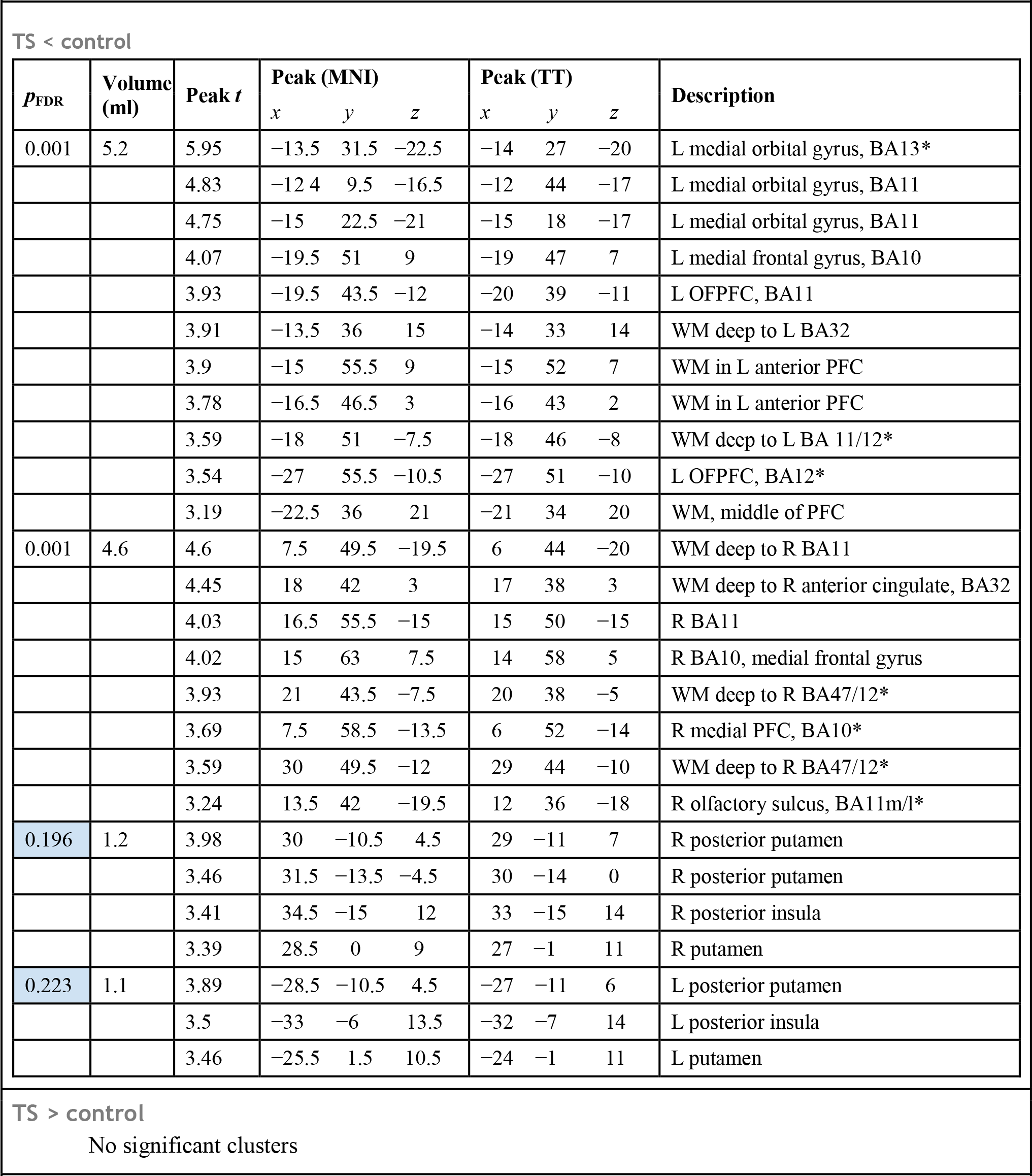
VBM results: white matter. **Legend:** *p*_roR_ = False discovery rate corrected*p* value for a suprathreshold cluster of this size in the *t* image. For each local maximum (peak) in the cluster, the table lists the *t* statistic at that voxel (193 df) and the atlas coordinates of that voxel's location. MNI = Montreal Neurological Institute template brain coordinates. TT = Talairach and Tournoux atlas coordinates. L = left hemisphere; R = right hemisphere. PFC = prefrontal cortex. OFPFC = orbitofrontal prefrontal cortex. BA = Brodmann area. * = description taken from (Öngür and Price, 2000).

**Supplemental Figure 1.**
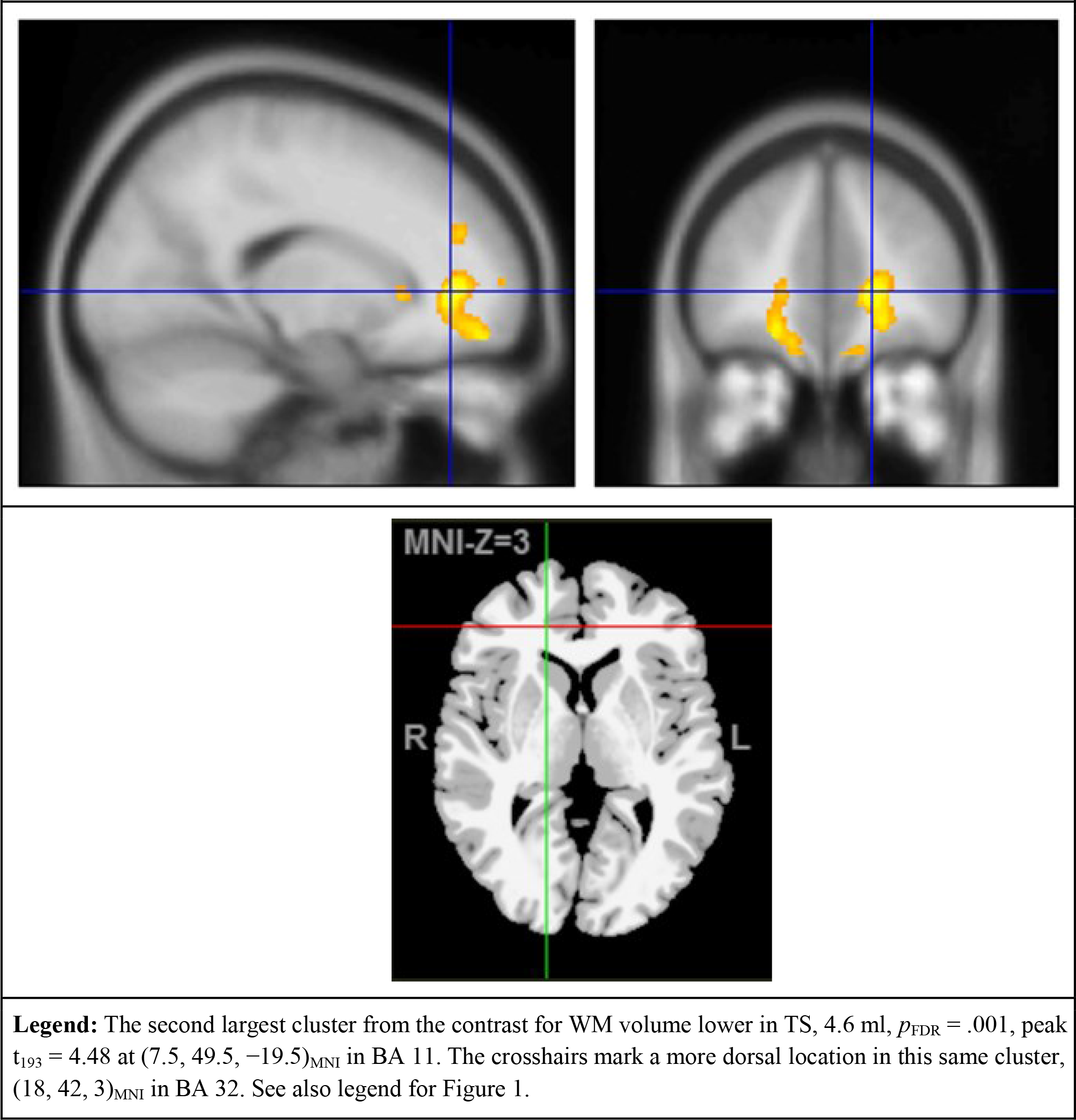
Second largest cluster showing lower WM volume in TS compared to controls **Legend:** The second largest cluster from the contrast for WM volume lower in TS, 4.6 ml, *p*_FDR_ = .001, peak *t*_193_ = 4.48 at (7.5, 49.5, −19.5)_MNI_ in BA 11. The crosshairs mark a more dorsal location in this same cluster, (18, 42, 3)_MNI_ in BA 32. See also legend for Figure 1.

Two additional symmetric clusters of decreased WM volume are of interest, though they did not remain statistically significant after multiple comparisons correction (each p = 0.2). These clusters include parts of posterior putamen and insula bilaterally (see Supplemental Figure 2).

**Supplemental Figure 2.**
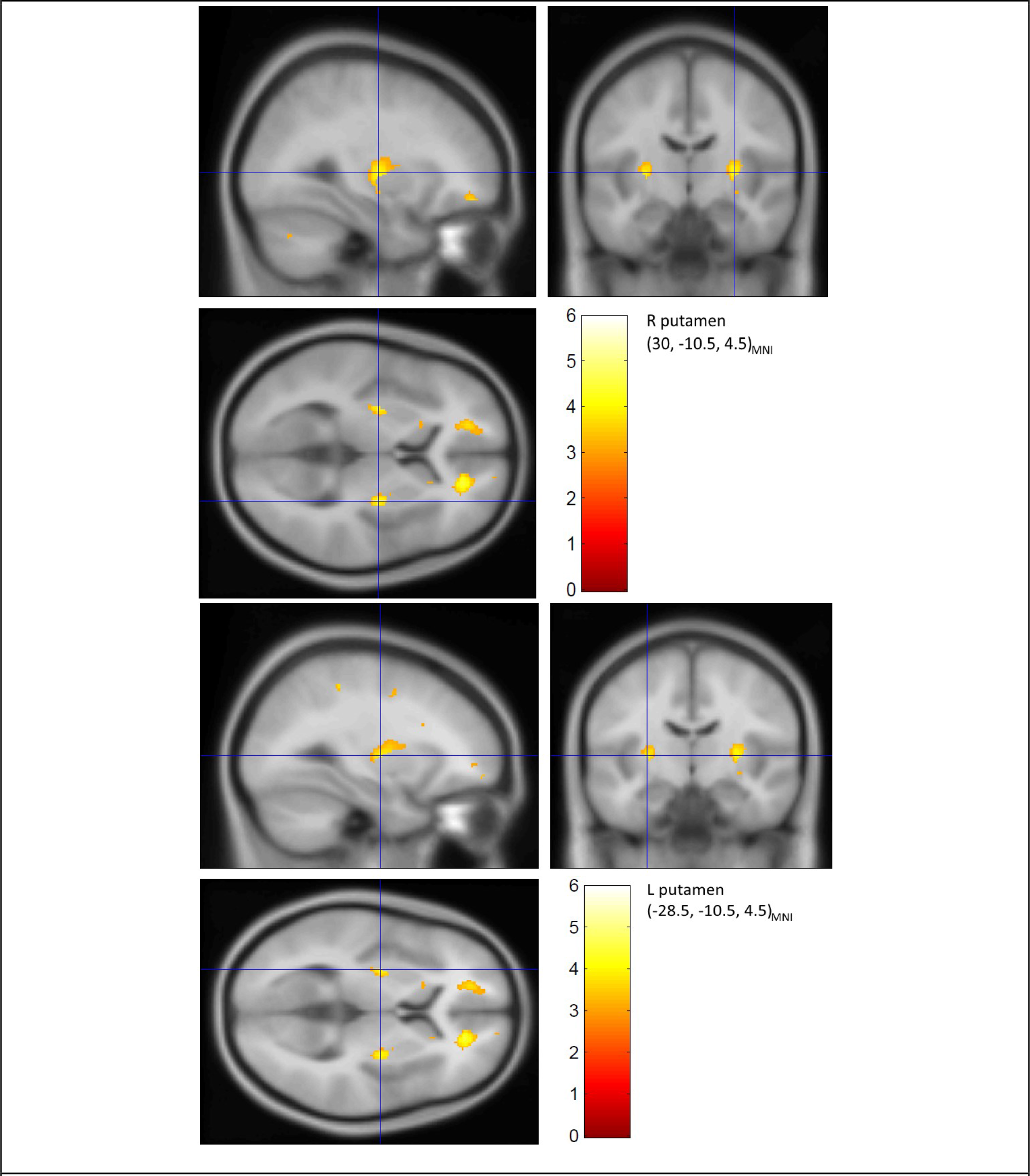
Additional clusters showing lower WM volume in TS compared to controls **Legend:** Clusters 3 and 4 from the contrast showing WM volume lower in TS than in the control group. The right peak is *t* = 3.98 at (30, −10.5, 4.5)MNI = (29, −11, 7)_TT_, 1.2ml, corrected *p* = 0.20, and the left peak is *t* = 3.89 at (−28.5, −10.5, 4.5)mn = (−27, −11, 6)_TT_, 1.1ml, corrected *p* = 0.22. See also legend to Figure 1.

### >Regional differences in gray matter volume in TS

Two clusters showed statistically significant increased GM volume in TS after correction for multiple comparisons (Table 3). The largest suprathreshold cluster had a volume of 4.4ml (corrected *p* = .001), with the peak *t* value (4.62, 193 d.f.) in the pulvinar nucleus of the left thalamus (see Figure 2a). A homologous cluster in the right pulvinar, peak *t* = 4.13 at (16.5, −28.5, −4.5)_MNI_ = (15, −29, 0)_TT_, was 1.6 ml, below the significance threshold (corrected *p* = 0.070). The second largest cluster had volume 2.7ml (corrected *p* = 0.011), with peak *t* = 4.06, and included the hypothalamus bilaterally and the ventral midbrain (see Figure 2b).

**Table 3.**
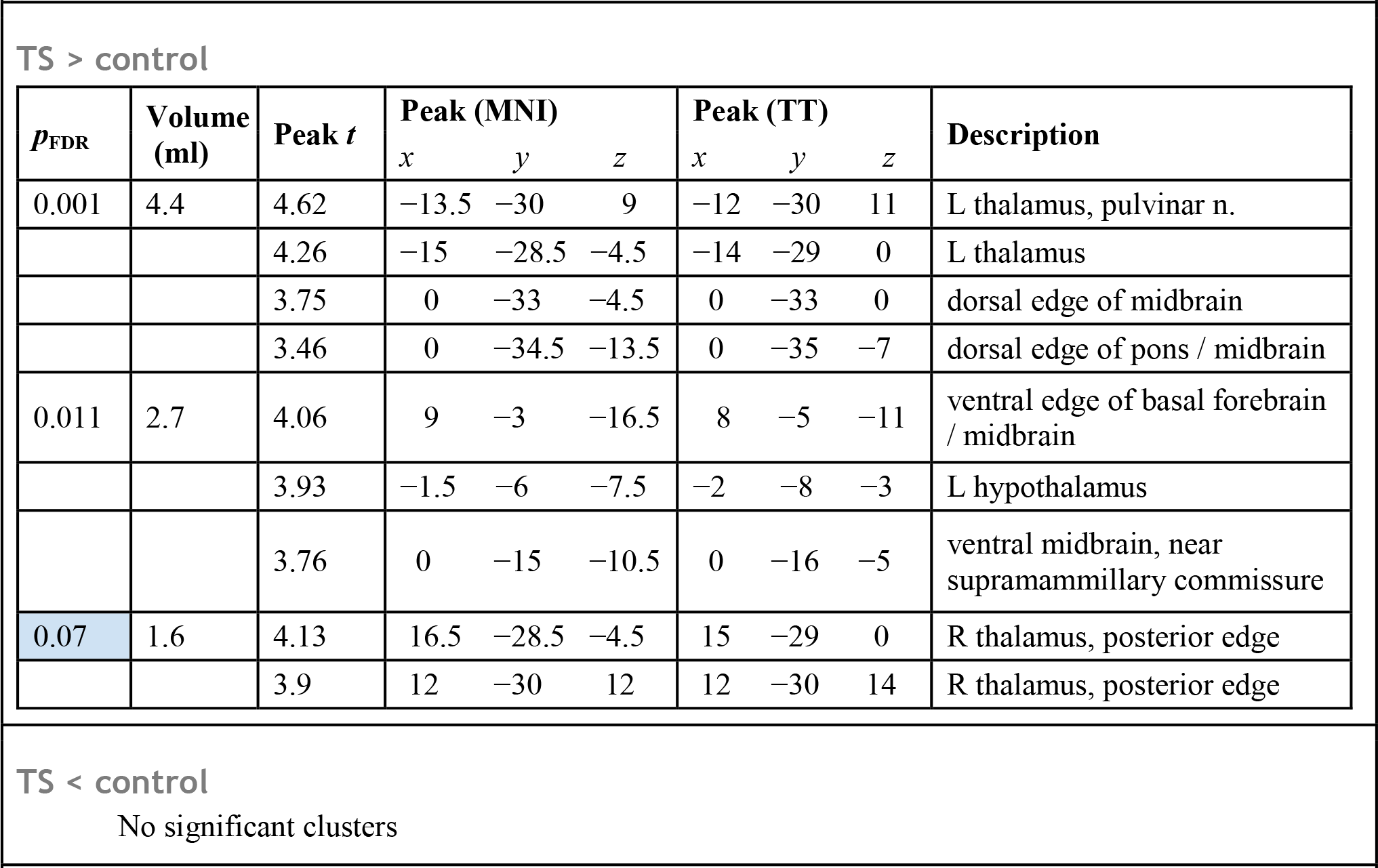
VBM results: gray matter. **Legend:** See legend for Table 2.

TS > control
| $p_{\text{FDR}}$ | Volume (ml) | Peak $t$ | Peak (MNI) | | | Peak (TT) | | | Description |
| --- | --- | --- | --- | --- | --- | --- | --- | --- | --- |
| | | | $x$ | $y$ | $z$ | $x$ | $y$ | $z$ | |
| 0.001 | 4.4 | 4.62 | -13.5 | -30 | 9 | -12 | -30 | 11 | L thalamus, pulvinar n. |
|  |  | 4.26 | -15 | -28.5 | -4.5 | -14 | -29 | 0 | L thalamus |
|  |  | 3.75 | 0 | -33 | -4.5 | 0 | -33 | 0 | dorsal edge of midbrain |
|  |  | 3.46 | 0 | -34.5 | -13.5 | 0 | -35 | -7 | dorsal edge of pons / midbrain |
| 0.011 | 2.7 | 4.06 | 9 | -3 | -16.5 | 8 | -5 | -11 | ventral edge of basal forebrain / midbrain |
|  |  | 3.93 | -1.5 | -6 | -7.5 | -2 | -8 | -3 | L hypothalamus |
|  |  | 3.76 | 0 | -15 | -10.5 | 0 | -16 | -5 | ventral midbrain, near supramammillary commissure |
| 0.07 | 1.6 | 4.13 | 16.5 | -28.5 | -4.5 | 15 | -29 | 0 | R thalamus, posterior edge |
|  |  | 3.9 | 12 | -30 | 12 | 12 | -30 | 14 | R thalamus, posterior edge |

**Figure 2.**
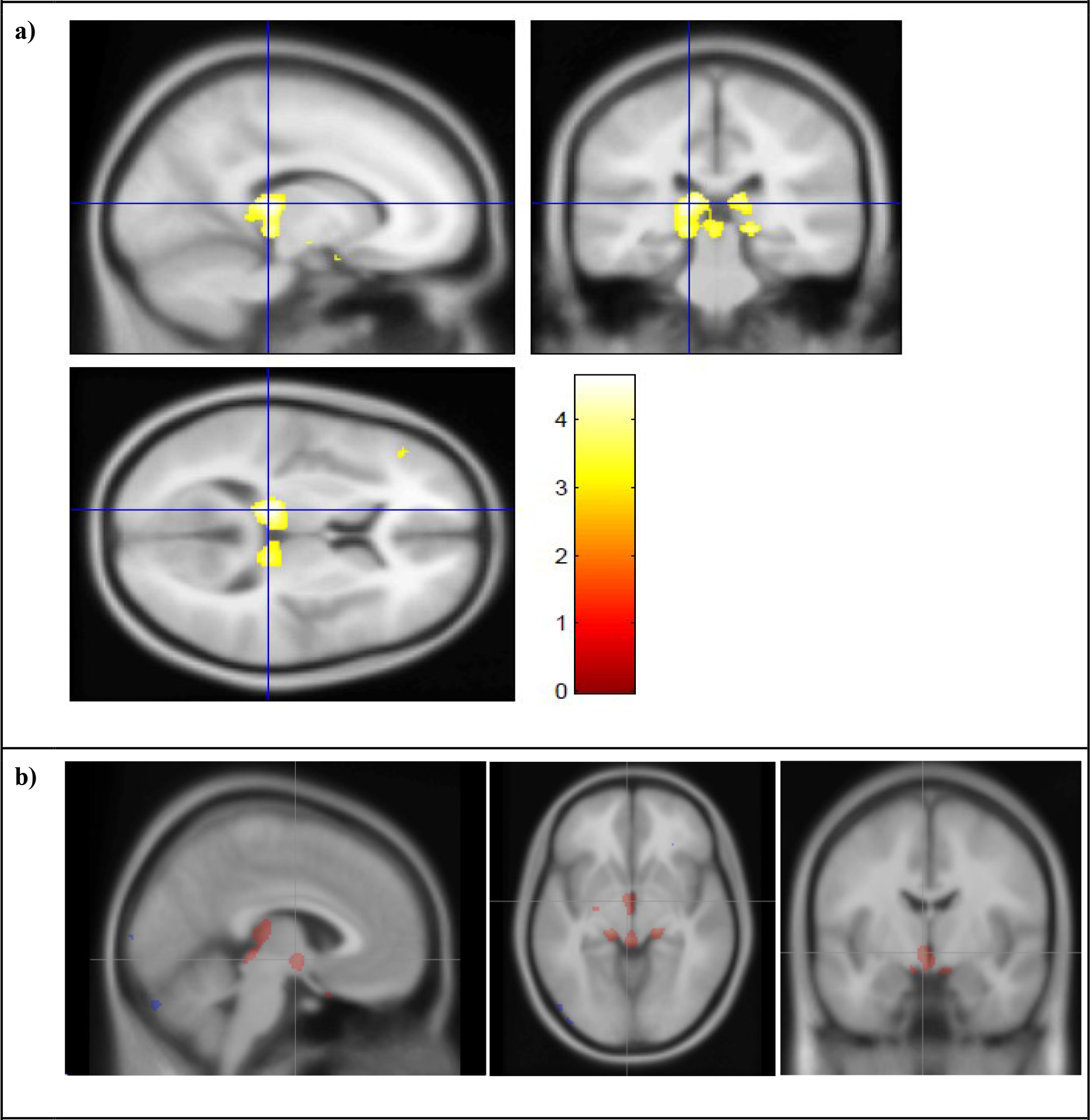
Largest Clusters Showing Greater GM Volume in TS Compared to Controls **Legend:** Largest cluster from GM > control contrast, in left pulvinar nucleus of thalamus (see Table 3 and legend to Fig. 1). **b)** From the second largest cluster from the GM > control contrast, with the crosshairs at (4, 6, −6)_MNI_ in hypothalamus. In this figure, all voxels with *t* ≥ 3.0 are highlighted in color to better visualize the underlying anatomy.

### Secondary analysis: scanner and MR sequence

The statistical model included a factor to account for different scanners or MR sequences, but such statistical control may be imperfect. Accordingly, we checked whether the findings from the overall group would still be present if the different-scanner concern were eliminated. One site acquired images from 46TS subjects and 27 control subjects on one scanner using the same sequence. For the left thalamus SPM cluster, for instance, the question is whether GM volume was higher in TS, as it was in the overall analysis, in these subjects who were all scanned on the same scanner with the same MR sequence. This question was tested using ANCOVAs with relative GM volume in the SPM cluster as the dependent variable, diagnosis and sex as factors, age as a covariate, and interactions of sex with diagnosis and age.
As in the full SPM analysis, this cluster was larger in TS (**Figure 3a**, diagnosis factor p=.004). Similarly in this same-sequence subgroup, the hypothalamus GM cluster was larger in TS (**Figure 3b**, diagnosis p=.000, with a significant diagnosis by sex interaction p=.009), and the OFPFC WM clusters were smaller in TS (**Figure 3c,d**, left WM diagnosis p=.000, right WM p=.000).

### Secondary analysis: IQ and comorbidity

We did not have IQ estimates from all sources, but we did for all but 1 subject from the single-sequence group discussed in the previous section. In this group, IQ differed between the two groups (TS 107.5 ± 11.9, control 117.8 ± 13.1, *p*<.002, unpaired *t* test), so we checked whether IQ explained any of the primary group differences by modeling relative cluster volume in each subject by ANCOVA with sex as a factor, age and IQ as covariates, and all interactions. Neither IQ nor interactions with IQ were significant for any of the 4 significant clusters (*p* for IQ was .08 for R OFPFC WM, .27 for L OFPFC WM, .10 for L thalamus, and .51 for hypothalamus).

ADHD was recorded for all TS subjects in that same subgroup. Using a similar approach, with sex and ADHD diagnosis as factors, age as a covariate, and all interactions, neither ADHD nor interactions with ADHD were significant for any of the 4 clusters. OCD diagnosis was not recorded in this subgroup, but as a loose proxy we dichotomized TS subjects based on OCD symptom severity (CY-BOCS scores, zero *vs.* greater than zero). This OCD factor was not significant, nor were any interactions with this factor, using the same analysis strategy as for ADHD.

**Figure 3.**
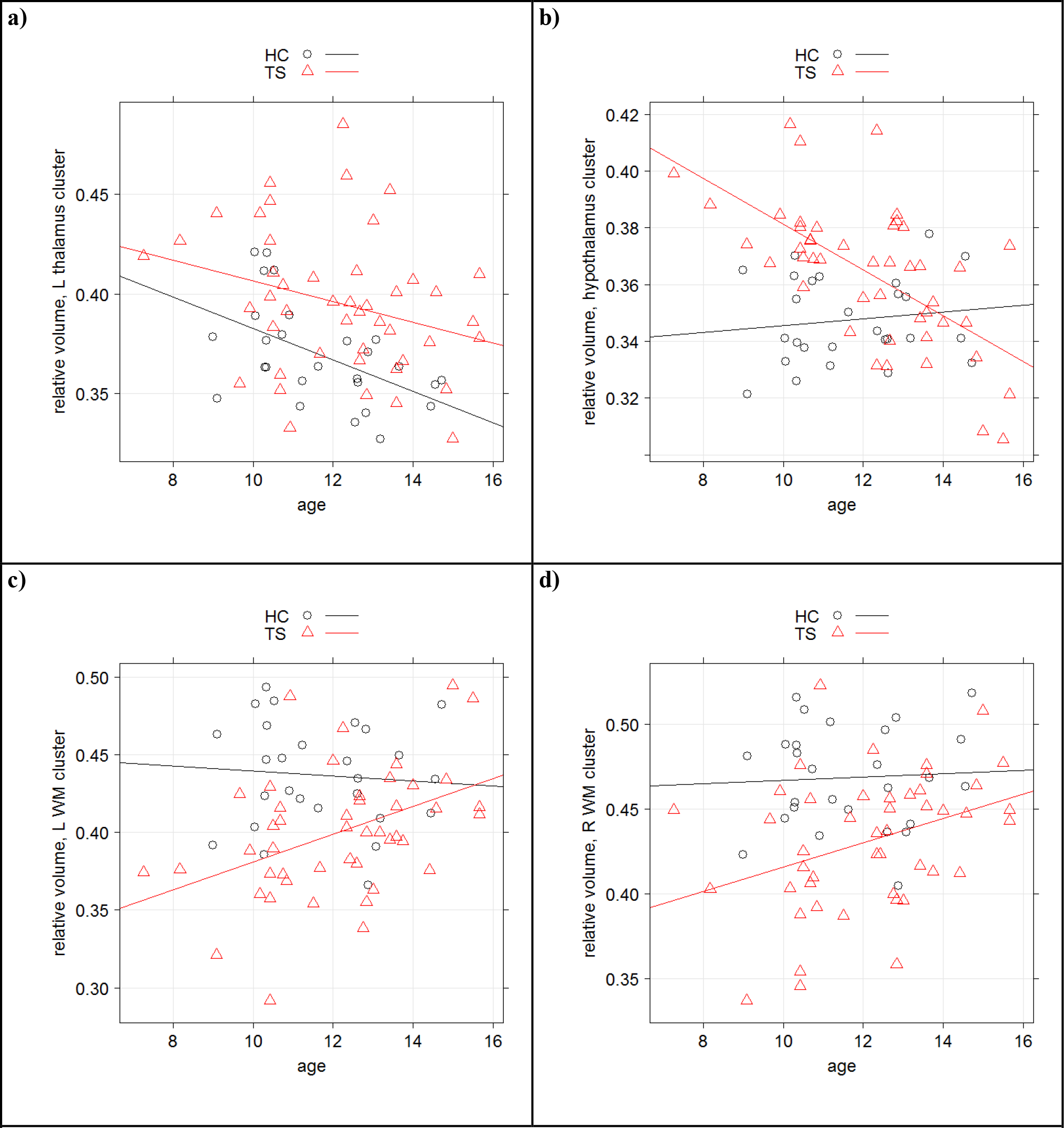
Key results hold in the single-site, same-sequence subgroup **a)** Relative GM volume of L thalamus cluster in a subgroup of subjects studied at one institution using a single MR sequence, plotted against age for TS subjects (triangles and red regression line) and control subjects (circles and black line). **b)** Same plot for hypothalamus. **c)** Same plot for relative volume of L OFPFC WM cluster. **d)** Same plot for R OFPFC WM cluster.

## Discussion

Here we present the largest study of brain structure ever reported in children with TS. We matched control subjects strictly for age, sex and handedness, and the statistical analysis used conservative methods to minimize Type I error. Our main findings were that the TS group had lower white matter volume than the control group deep to orbital and medial prefrontal cortex, and greater gray matter volume in the posterior thalamus and hypothalamus.

### Lower WM volume in prefrontal cortex in TS

The finding of decreased WM in orbital cortex is consistent with the decreased volume found in a predefined orbital frontal cortex area in an earlier large structural MRI study in TS (Peterson et al., 2001). The present results suggest that the decreased orbital prefrontal cortex (OFPFC) volume may be attributable to WM. Another study focusing specifically on white matter (Cheng et al., 2014) identified 10 WM tracts with decreased WM integrity (scaled fractional anisotropy) in a group of 15 currently unmedicated adults with TS compared to 15 tic-free control subjects matched for age and sex. Four of those 10 tracts involved the OFPFC, specifically those connecting OFPFC with pre-SMA (anterior to supplementary motor cortex), ventral premotor cortex, primary motor cortex, and supplementary motor cortex.

The orbital prefrontal cortex regions identified in the present study have been linked to behavioral disinhibition and to visceral and emotional responses (Knutson et al., 2015; Ongur and Price, 2000; Barbas et al., 2003). Thus, reduced WM volume in this region may contribute to the inability to inhibit unwanted behaviors, namely tics. Alternatively, the connection of OFPFC to tics may be sensory. Most tic patients report that tics are often responses to uncomfortable internal sensations, like a tickle in the throat before a cough. Many tic experts, though not all, conclude that these premonitory sensations may be the primary phenomenon rather than the observed tics (Leckman et al., 2013). Abnormal white matter connections to ventromedial OFPFC, a brain region involved in assessing internal sensations, fit well with such a model.

The clusters of decreased WM volume also extended to pregenual WM and WM deep to medial frontal gyrus (BA 10). A previous study that examined WM integrity in TS found a strong correlation between past-week tic severity and WM fractional anisotropy deep to superior frontal gyrus (Müller-Vahl et al., 2014). Increased tic severity was associated with decreased WM integrity in this region, consistent with our results, though the specific location of the region was 10mm superior to the BA 10 peak in the current study.

#### Putamen

The paired clusters of decreased WM volume in posterior putamen are interesting given the posterior putamen's prominent role in movement. Left and right putamen were the locations at which apparent diffusion coefficient was most highly correlated with tic severity (Müller-Vahl et al., 2014), though the most significant voxels in that study are 16-17 mm anterior and inferior from the peaks reported here. However, the clusters in the present study were not significant after correction for multiple corrections, and a WM difference might be more easily interpreted as referring to the external capsule or extreme capsule than the putamen itself. These clusters also extended to bilateral insula.

#### Other

Some previous TS studies found larger volume or reduced fractional anisotropy in corpus callosum, but otherwise, previous WM findings in TS have been variable (Greene et al., 2013).

### Greater GM volume in pulvinar nucleus, midbrain and hypothalamus in TS

#### Pulvinar

Several imaging studies have examined thalamic volume in TS (Greene et al., 2013). The largest of these found increased total thalamic volume in children and adults with TS (about 5%), with outward deformation (bulges) compared to thalamic shape in control subjects (Miller et al., 2010). The most prominent differences were found on the ventral, lateral and posterior surfaces, corresponding to several motor nuclei and the pulvinar. Thus, that study's results are quite consistent with the present finding of greater GM volume in the pulvinar in a large group of children and adolescents. Miller *et al.* posit several possible explanations for finding enlargement in these thalamic regions, including hyperactive motor circuitry, compensatory mechanisms derived from years of controlling (or attempting to control) tics, or secondary GM changes in the face of WM alterations.

The medial pulvinar nucleus is widely connected to cortex, including prefrontal, orbital, and cingulate cortical areas; the lateral pulvinar projects to parietal, temporal, and extrastriate regions; and the inferior pulvinar has bidirectional connections with visual cortical regions (Benarroch, 2015; Shipp, 2003). Given these widespread projections and innervations, we speculate that increased GM volume in TS may relate to multisensory integration in the thalamus, or to the linking of sensory input to cognitive-, motivational-and movement-related areas of cortex. Sensory symptoms are common in TS (premonitory urges, hypersensitivity), yet TS patients do not demonstrate primary sensory deficits (Belluscio et al., 2011; Schunke et al., 2016), suggesting involvement of higher-order functions such as attention. There is evidence for a functional role of the pulvinar nucleus in spatial attention and attention to salient stimuli (Grieve et al., 2000; Shipp, 2004).

#### Midbrain

Part of the thalamus GM cluster includes dorsal midbrain. Interestingly, a VBM study of 31 patients and 31 controls also identified a significant increase in GM volume in midbrain (Garraux et al., 2006), though that statistical peak was inferior and anterior to the one identified in the present study.

#### Hypothalamus

One cluster of increased GM volume included hypothalamus. We are not aware of any previous studies linking TS to this structure. However, the hypothalamus does receive inhibitory innervation from the ventral medial OFPFC via the central nucleus of the amygdala (Ongur and Price, 2000; Barbas et al., 2003) . This anatomical connection is intriguing given the OFPFC WM changes in this study. Future work may study the hypothalamus more specifically.

### Comparison to OCD

The main findings in the present study largely overlap with previous structural MRI results in OCD. OCD is relevant here for several reasons, including the high comorbidity rates of OCD and TS and the phenomenological similarities in symptoms between these conditions, as thoroughly reviewed recently by (Eddy and Cavanna, 2014). The most prominent overlapping finding between previous OCD studies and the present TS study is volumetric differences in the orbitofrontal cortex in OCD. However, these reports are mixed, including both decreased orbitofrontal volume (Atmaca et al., 2007), increased orbitofrontal volumes (Valente et al., 2005; Kim et al., 2001) thicker left and thinner right orbitofrontal cortex related to an increased likelihood of responding well to treatment (Hoexter et al., 2015). Similarly, left and right OFPFC GM volume correlated in opposite directions with symmetry/ordering symptom severity in OCD (Valente et al., 2005). The potential importance of these structural results is highlighted by functional imaging studies; a meta-analysis of PET and SPECT studies in OCD found that the largest effect sizes for abnormal function in OCD were for left (d = 1.15) and right (d = 1.04) orbital gyrus (Whiteside et al., 2004).

Also consistent with previous reports in OCD is the present finding of increased GM volume in thalamus and hypothalamus. Larger thalamic volumes have been reported in OCD using volume-of-interest (Atmaca et al., 2007) and VBM approaches (Kim et al., 2001). Kim et al. also found increased GM volume in bilateral hypothalamus in OCD.

Still, OCD *per se* is unlikely to explain our current results, given null results in the secondary analysis based on current OCD severity. However, this conclusion should be confirmed in a sample with prospective, systematic psychiatric diagnosis.

### The dog that did not bark in the night …

A word is due about previous volumetric findings that were not replicated here. The most notable of these is decreased caudate volume in TS reported by Peterson et al (Peterson et al., 2003) in a study of 154 children and adults with TS and 130 controls (including a total of 173 children), and by two other groups (Makki et al., 2008; Peterson et al., 1993; Müller-Vahl et al., 2014; Makki et al., 2009; Müller-Vahl et al., 2009). The potential relevance of that finding was enhanced by the report that smaller caudate volume in childhood predicted worse tic severity in young adulthood, showing that decreased caudate volume could not be just a consequence or adaptation of the brain to tics (Bloch et al., 2005). Conceivably our caudate non-finding reflects a Type II error.

On the other hand, several other studies did not find a significantly smaller caudate in TS (Black et al., 2003; Moriarty et al., 1997; Zimmerman et al., 2000; Garraux et al., 2006; Kassubek et al., 2006; Ludolph et al., 2006; Williams et al., 2013; Wang et al., 2007; Roessner et al., 2011; Wittfoth et al., 2012; Roessner et al., 2009). The largest of these included 49 boys with TS and 42 boys without tics (Roessner et al., 2011). The present study has some merits relative to previous studies beyond the larger sample size; it excluded adults and handled variation in age and sex by one-to-one age matching in addition to statistical accounting for linear effects of age. Furthermore, the caudate is a relatively small structure, surrounded by white matter and CSF, and hence especially susceptible to partial volume effect and, presumably, to the artifactual reduction in volume with frequent small-amplitude head movements demonstrated with other techniques (Alexander-Bloch et al., 2016; Reuter et al., 2015). Therefore caudate volume may be normal in TS.

### Limitations

The most important limitation is the use of different scanners, different sequences, and possibly different recruitment sources or diagnostic methods across sites. However, the analysis of the largest subgroup suggests strongly that the key findings are not driven by differences in site, scanner or sequence.

A second limitation is that phenotypic data are limited for many of the “legacy” subjects. For instance, diagnosis (TS vs. chronic motor tic disorder) is missing for some subjects in the TS group, and for many subjects we have limited information on comorbid diagnosis. In the available data our key findings are not significantly linked to IQ, ADHD or OCD, but of course future studies will benefit from adequate prospective assessment of all these variables (Castellanos et al., 1996).

Recently, small head movement not detected by visual inspection of MR images has been shown to artifactually lower GM volume in VBM analyses, presumably by a mechanism similar to partial volume effect (Alexander-Bloch et al., 2016; Reuter et al., 2015). Fortunately, group differences in residual head movement cannot easily explain the decreased WM volume or the increased GM identified in this study, since the TS group would be expected to show more head movement.

### Future directions

A prospective study design with additional clinical information can test whether the posterior thalamic finding in fact relates to sensory symptoms in TS, whether the OFPFC finding relates to measures of response inhibition, coprophenomena (Black et al., 2014; Eapen et al., 2015), or socially inappropriate non-tic behavior (Kurlan et al., 1996), and to what extent the severity of tics, obsessions and compulsions explain these findings. Future structural imaging studies can help elucidate at what age the regional differences in GM and WM volume in TS first manifest and whether they persist into adult life, thus helping to clarify whether the volumetric differences represent failures of maturation or alterations after a period of normal development.

Studies with different methodology will be required to elucidate the mechanism responsible for the volumetric abnormalities. Postmortem studies in TS have not typically focused on the regions identified in this study (Swerdlow and Young, 2001). Thus it is not clear whether, for example, increased GM volume in posterior thalamus reflects increased neuronal cell number, glial cell number, neuropil (*e.g.* deficient pruning), or increased water content. On the other hand, this reflects a potential strength of the present study: an unbiased, whole-brain analysis identified regions of brain that have hardly been studied in TS.

## Methods

This study was approved by the Washington University Human Research Protection Office (IRB), protocol # 201108220. Most MR images and clinical information were originally collected under different IRB protocols (independent of this study) at the 4 imaging sites: WUSM, NYU, KKI, UCLA. Herein we call this “legacy” data. The WUSM, KKI and UCLA sites enrolled additional new subjects specifically for this study. The transmission of any human subjects data to the consortium was approved by each site's respective IRB. Some data were provided anonymously to the consortium under code-sharing agreements.

Imaging data were stripped of personal identifiers such as name and date of birth and archived at the Central Neuroimaging Data Archive (CNDA) hosted at http://cnda.wustl.edu (Gurney et al., 2015). REDCap electronic data capture tools hosted at Washington University were used to manage the clinical data collected at WUSM (Harris et al., 2009).

## Subjects

T1-weighted MPRAGE images were located from 230 children aged 7–17 with DSM-5 TS or chronic ticdisorder and 216 tic-free controls. Of the subjects with at least one remaining image, 103 TS and 103 controls could be matched 1:1 for age (within 0.5 year), sex, and handedness (Figure 4).

MPRAGE (3D T1-weighted) data were acquired on several MR scanners with varying parameters. The most common structural image protocol was an MP-RAGE with total scanning time 6-10 minutes and voxel size 1.0-1.25mm^3^ (Supplemental Table 1). Authors ACW, DJG or KJB visually reviewed each structural MRI and excluded images with any visible artifact in the brain. The provenance of the images used in this analysis is described further in Supplemental Table 1.

**Figure 4.**
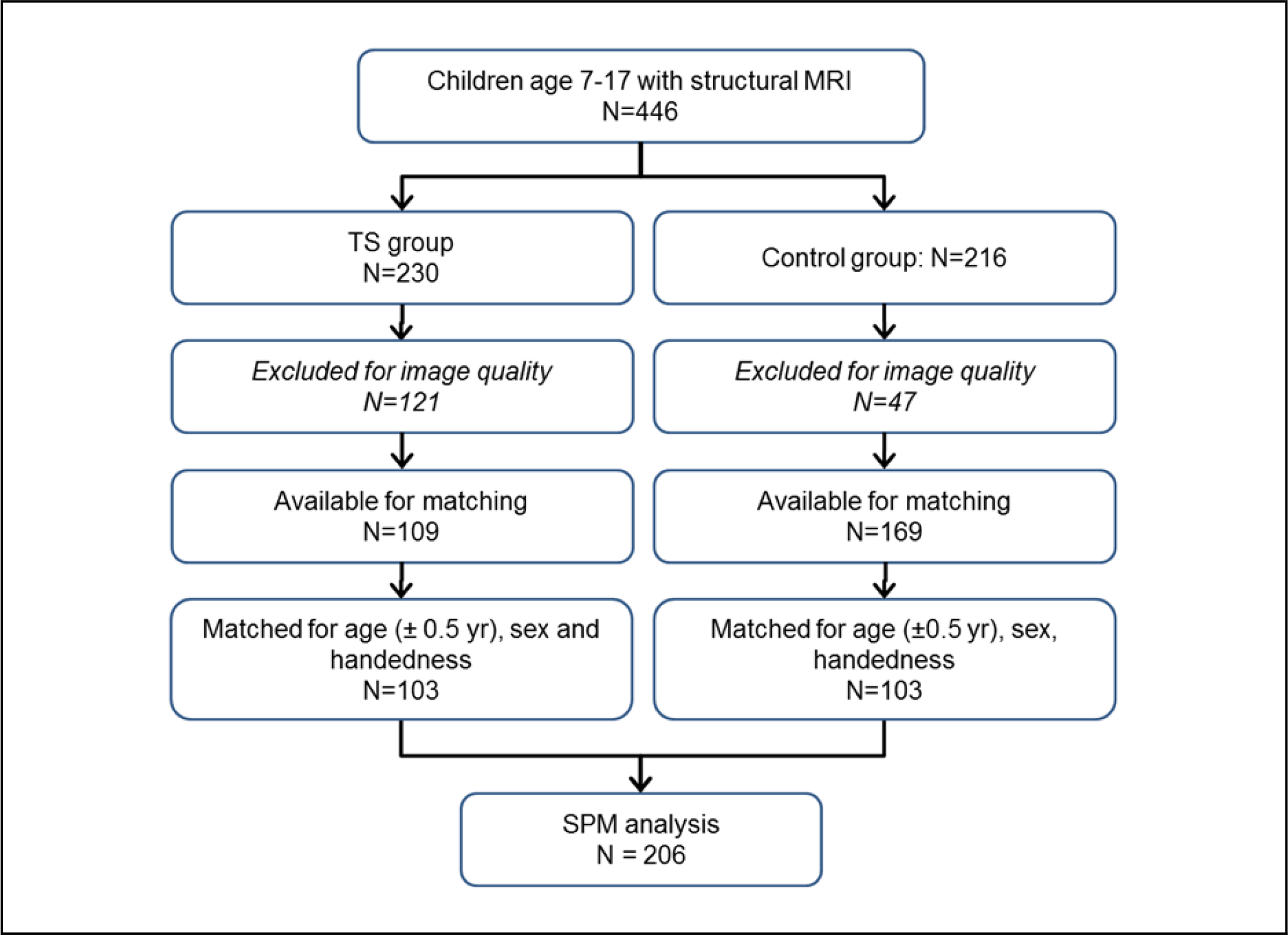
Subject flow diagram

## Image processing

If a subject had more than one MPRAGE image of adequate quality, these images were averaged after mutual rigid-body alignment using a validated method (Black et al., 2001), and the mean imagerepresented that subject in all subsequent steps. Beyond this point all image analyses were performed with SPM software v.12b using the method of J. Ashburner (Mechelli et al., 2005; Perantie et al., 2011).

Each subject's image was nonlinearly normalized to MNI space, and the atlas-aligned images were averaged to create an MPRAGE template specific to this study, as described elsewhere (http://irc.cchmc.org/software/tom.php) (Wilke et al., 2008). For each subject, segmented images were created to reflect the probability that each voxel was composed of gray matter (GM), white matter (WM), or CSF. This computation used a Bayesian approach, with prior probabilities established by population templates for GM, WM and CSF to inform interpretation of the subject's MR signal at each voxel. Alignment and segmentation were then refined by tissue-specific realignment (Good et al., 2001). The tissue density images were multiplied by the local volume in the subject image corresponding to each voxel in the atlas template to produce images showing at each atlas voxel that subject's GM, WM and CSF volume contributing to that voxel. The GM volume image from one subject is shown in Supplemental Figure 3. A 3D Gaussian filter (FWHM 6mm) was applied to the GM and WM images and the smoothed images were submitted to SPM analysis.

## Analysis

Statistical image analysis was performed using SPM software v. 12b (http://www.fil.ion.ucl.ac.uk/spm/software/), which computed at each voxel a general linear model (GLM) with dependent variable gray matter volume (GM), factors diagnostic group, MRI scanner and sequence (Supplemental Table 1), and sex; age at scan as a covariate; and interactions of group × sex and sex × age. Proportional scaling by each subject's total GM + WM volume corrected for global brain volume. The GM analysis was limited to voxels at which GM concentration was >20%. The WM analysis used the same methods.

One-tailed contrasts were used to generate t images comparing TS and control groups. Statistical significance was determined by the volume of clusters defined by contiguous voxels with ∣t∣>3.0, corrected to a false discovery rate of 5%. Peak voxel locations in MNI space were transformed to Talairach atlas coordinates using MNI2TAL (http://bioimagesuite.yale.edu/mni2tal) (Lacadie et al., 2008).

**Supplemental Figure 3.**
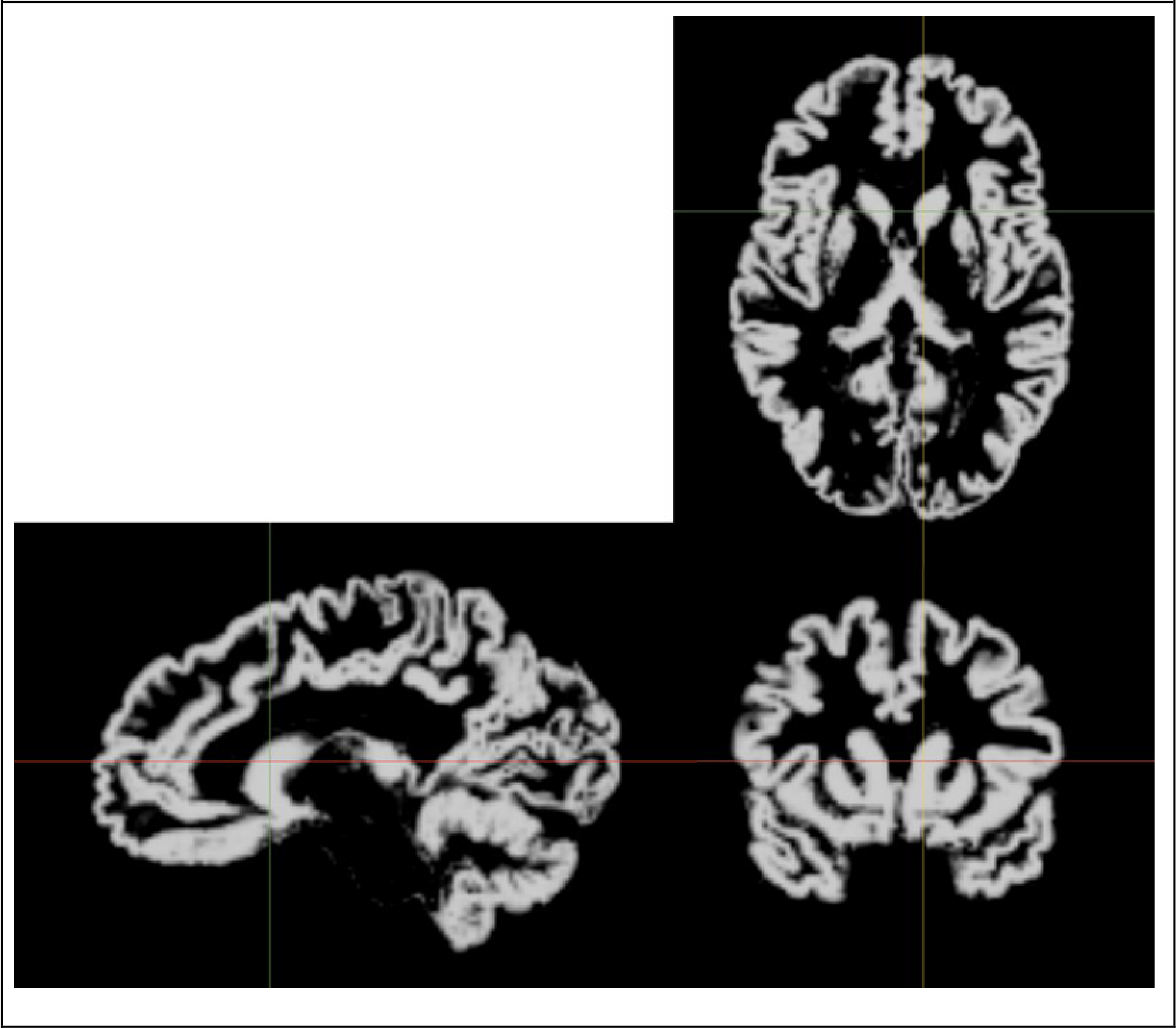
GM volume image from one subject

Total GM and total WM were modeled similarly, *i.e.* with diagnostic group, MRI sequence, and sex as factors, age at scan as a covariate, and interactions of group × sex and sex × age, but of course omitting the global volume correction, using R statistical software v. 3.1.2 (R Core Team, 2015; RStudio Team, 2014).

Secondary analyses focused on the key findings from the SPM analyses. For each significant cluster from the SPM analyses of GM, the sum of each subject's GM volume over all voxels in that cluster was corrected for the subject's total brain volume (GM + WM) by division. The same was done for the significant WM clusters. These relative cluster volumes for each subject were the dependent variables to test for effects of scanner and sequence, ADHD, and IQ, either in the entire TS group or in the subjects scanned with sequence 3 from Supplemental Table 1.

## Authorship note

Consortium investigators, collaborators and other study personnel include:

Bradley L. Schlaggar, Investigator, WUSTL,* consortium co-chair; Kevin J. Black, Investigator, WUSTL, consortium co-chair; Deanna J. Greene, Investigator, WUSTL; Jessica A. Church, Collaborator, WUSTL, current affiliation University of Texas at Austin; Steven E. Petersen, Collaborator, WUSTL; Tamara Hershey, Collaborator, WUSTL; Deanna M. Barch, Collaborator, WUSTL; Joan L. Luby, Collaborator, WUSTL; Alton C. Williams, III, Study staff, WUSTL, medical student researcher, current affiliation MUSC; Jonathan M. Koller, Study staff, WUSTL, senior statistical data analyst; Matthew T. Perry, Study staff, WUSTL, postdoctoral research rotation; Emily C. Bihun, Study staff, WUSTL, site coordinator and psychiatric interviewer; Samantha A. Ranck, Study staff, WUSTL, psychiatric interviewer; Adriana Di Martino, Investigator, NYU; F. Xavier Castellanos, Investigator, NYU; Barbara J. Coffey, Investigator, NYU and Icahn Mount Sinai School of Medicine; Michael P. Milham, Investigator, NYU, current affiliation Child Mind Institute, New York, NY; Abigail Mengers, Study staff, NYU; Krishna Somandepalli, Study staff, NYU; Stewart H. Mostofsky, Investigator, KKI and JHU; Harvey S. Singer, Investigator, JHU; Carrie Nettles, Study staff, KKI; Daniel Peterson, Study staff, KKI; John Piacentini, Investigator, UCLA; Susan Y. Bookheimer, Investigator, UCLA; James T. McCracken, Investigator, UCLA; Susanna Chang, Investigator, UCLA; Adriana Galván, Investigator, UCLA; Kevin J. Terashima, Study staff, UCLA; Elizabeth R. Sowell, Investigator, University of Southern California and Children's Hospital Los Angeles.

* WUSTL = Washington University in St. Louis. NYU = New York University. KKI = Kennedy Krieger Institute. JHU = Johns Hopkins University. UCLA = University of California, Los Angeles.

## Acknowledgments

We thank Drs. Carol A. Mathews, Keith A. Coffman, Jeremy D. Schmahmann and Barry S. Fogel for helpful comments on the Discussion.

We gratefully acknowledge funding by the Tourette Association of America and its donors, by the U.S. National Institutes of Health (NIH; grants K24 MH087913, P30 CA091842, P50 MH077248, UL1 TR000448, K01 MH104592, and R21 NS091635), by the Brain & Behavior Research Foundation (NARSAD Young Investigator Award) and by the Siteman Comprehensive Cancer Center. Research reported in this publication was also supported by the Eunice Kennedy Shriver National Institute Of Child Health & Human Development of the National Institutes of Health under Award Number U54 HD087011 to the Intellectual and Developmental Disabilities Research Center at Washington University. The content is solely the responsibility of the authors and does not necessarily represent the official view of the funders.

Key findings from this work were presented at the annual meeting of the American Neuropsychiatric Association, 27 March 2015 (Williams AC III, Greene DJ, Perry MT, Koller JM, Schlaggar BL, Black KJ, The Tourette Syndrome Association Neuroimaging Consortium: A multi-site voxel-based morphometry study of Tourette syndrome. J Neuropsychiatry Clin Neurosci 27 (2): e190, 2015, doi:10.1176/appi.neuropsych.272Abstracts), and at the First World Congress on Tourette Syndrome & Tic Disorders, 25 June 2015 (Williams AC III, Greene DJ, Perry MT, Koller JM, Schlaggar BL, Black KJ, The Tourette Syndrome Neuroimaging Consortium: A pilot multicenter study of brain structure in pediatric Tourette syndrome, http://eventmobi.com/tourette2015/documents/112809, archived by WebCite^®^ at http://www.webcitation.org/6gLnEvFBT.

## Conflict of interest

The authors declare no conflicts of interest.

